# Structural variants in human congenital heart disease disrupt distal genomic regulatory contacts of developmental genes

**DOI:** 10.64898/2026.02.28.708767

**Authors:** Jodi Lee, Jingshing Wu, Maureen Pittman, Zoe L. Grant, Shuzhen Kuang, Daniel Quiat, Sarah U. Morton, Geoff Fudenberg, Michela Traglia, Kelly A. Hayes, Pediatric Cardiac Genomics Consortium, Ritu Kumar, Benoit G. Bruneau, Katherine S. Pollard

**Affiliations:** Gladstone Institutes; San Francisco, CA, USA; TETRAD Graduate Program, University of California, San Francisco; San Francisco, CA, USA; Department of Pediatrics, University of Wisconsin School of Medicine and Public Health; Madison, WI, USA; Biomedical Informatics Graduate Program, University of California, San Francisco; San Francisco, CA, USA; Department of Pediatrics, Harvard Medical School and Boston Children’s Hospital; Boston, MA, USA; Department of Quantitative & Computational Biology, University of Southern California; Los Angeles, CA, USA; Department of Molecular Cell Biology, Harvard University; Cambridge, MA, USA; Roddenberry Center for Stem Cell Biology and Medicine at Gladstone, San Francisco, CA, USA; Department of Pediatrics, Cardiovascular Research Institute, Institute for Human Genetics, and the Eli and Edythe Broad Center for Regeneration Medicine and Stem Cell Research, University of California, San Francisco; San Francisco, CA, USA; Department of Epidemiology & Biostatistics, University of California, San Francisco; San Francisco, CA, USA; Chan Zuckerberg Biohub; San Francisco, CA, USA

## Abstract

Predicting the functional significance of structural variants (SVs) associated with genetic diseases remains challenging. To test the hypothesis that SVs from people with congenital heart disease (CHD) disrupt developmental chromatin interactions, we developed CardioAkita, a machine-learning model that predicts how variants alter 3D chromatin structure. Analyzing previously genotyped *de novo* SVs (*dn*SVs), we observed a positive association between CHD severity and CardioAkita scores across dozens of families. From whole-genome sequencing of three individuals with CHD we predicted disruptive *dn*SVs. Induced pluripotent stem cells engineered to harbor these variants confirmed CardioAkita’s predictions of 3D chromatin changes, and further revealed aberrant expression of local genes including cardiac developmental genes, suggesting that chromatin reorganization plays a significant mechanistic role in the genetic etiology of CHD. Our findings highlight the potential for models of 3D chromatin organization to predict the pathogenicity and underlying mechanisms of SVs in human disease.

## Introduction

Many human adult^1^ and pediatric^2^ diseases have a genetic basis, but variants of “undetermined significance” are common among individuals undergoing genetic testing^3,4^. These include structural variants (SVs), DNA changes such as deletions, inversions, insertions, or duplications that are among the most significant contributors to genetic variation in the human genome. Congenital heart disease (CHD) is the most common major congenital anomaly, presenting in about 1% of live births. SVs have been linked to CHDs and other developmental disorders, and their presence is typically associated with worse clinical outcomes^5–7^. Nonetheless, identifying the underlying genetic causes for many CHD cases remains challenging: 60% of sequenced individuals with CHD lack a known genetic etiology, because none of their variants are recognized as pathogenic. This diagnostic knowledge gap is particularly large for noncoding variants.

SVs can alter flanking chromatin interactions^8^, and in a few cases these effects have been linked to disease, including cancer, neurodegeneration, and cardiac defects^9–11^. Disruption of chromatin loops or domain boundaries can alter interactions between distal regulatory elements and promoters, leading to aberrant gene expression^12–14^. Since disruptive SVs are often distant from the affected genes, and experimentally measuring SV effects is not currently a scalable assay, it is challenging to predict whether an SV ascertained in a person with CHD is pathogenic and to predict if its molecular mechanism involves changes to 3D genome architecture.

Computational modeling provides a potential solution. Sequenced-based neural network models have enabled researchers to screen and prioritize genetic variants that likely alter transcription factor binding, chromatin states, or gene regulation^15–20^. We recently extended this approach to 3D genome organization by combining Akita, a model that accurately predicts chromatin interactions (Hi-C) from DNA sequence alone^21^, with the SuPreMo pipeline^22^ to score SVs from children with brain tumors^20^ or autism spectrum disorder^23^. Genome editing experiments have validated several of Akita’s SV predictions^21,23^, suggesting that this approach could be applied to causal variant discovery for CHD and other diseases. Major obstacles include making Akita’s predictions more cell type specific and expanding the model–previously trained using high-resolution Hi-C data from immortalized cell lines–to more disease relevant cellular contexts. In this study, we address these challenges with the objective of exploring the hypothesis that CHD-associated SVs disrupt chromatin interactions and gene regulation in heart development.

## Results

### CardioAkita predicts cardiomyocyte specific genome folding from DNA sequence

We leveraged high-resolution Hi-C data from WTC11 induced pluripotent stem cells (iPSCs) differentiating into atrial and ventricular cardiomyocytes^24^ to train a convolutional neural network to predict chromatin interaction maps along both time courses from the underlying DNA sequence. This CardioAkita model (**Fig. 1A**) has a similar architecture to Akita^25^, but the output head makes predictions for iPSCs, six atrial time points spanning cardiac mesoderm (D2, D4), cardiac precursors (D6), and cardiomyocytes (D11, D20, D45), and five ventricular time points spanning cardiac mesoderm (D2, D4), cardiac precursors (D6), and cardiomyocytes (D11, D23). The inclusion of a later atrial time point (D45) was based on bulk RNA-seq data indicating slower maturation^24^. The data preprocessing, loss, and partitioning of the genome into train/validation/test sets were identical to Original Akita (Methods).

**Figure 1.**
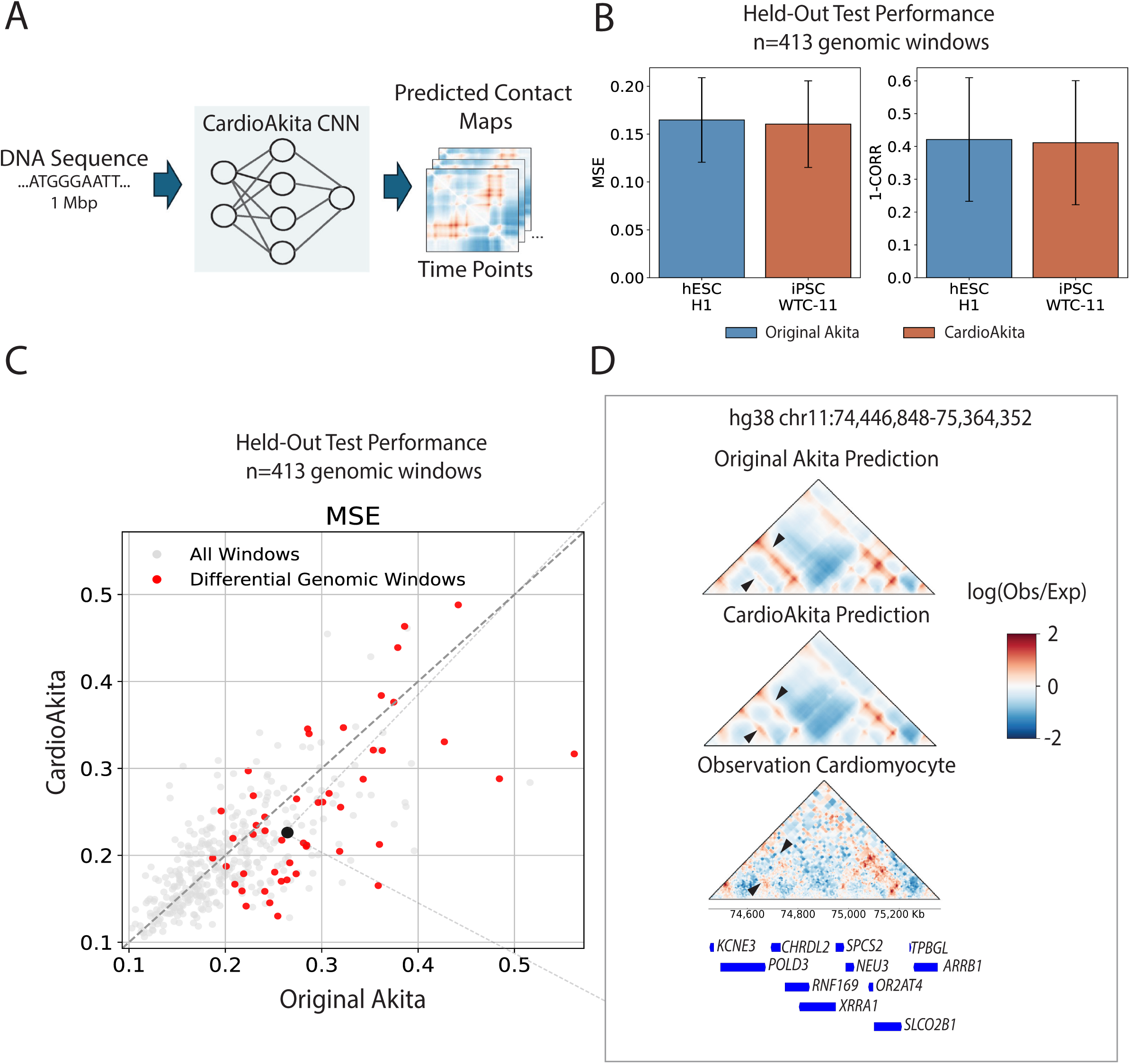
CardioAkita predicts 3D genome folding in differentiating cardiomyocytes from DNA sequence. **A.** Schematic of the CardioAkita model. From approximately 1-Mbp of 1-hot encoded DNA, we predict chromatin contact maps for time series of iPSCs differentiating into atrial and ventricular cardiomyocytes. **B.** Performance of Original Akita and CardioAkita on similar cell types, H1 hESCs (blue) and WTC11 iPSCs (red). MSE: mean squared error; 1-CORR: one minus Pearson correlation. Bar: mean performance across 413 held-out genomic windows; Error bars: +/- 1 standard deviation. **C.** On cardiomyocyte (ventricular D23) held-out loci, CardioAkita performs slightly better than Original Akita: 242/413 points below dotted line (59%). Only CardioAkita makes cardiomyocyte-specific predictions. Original Akita hESC predictions are shown here; other non-cardiomyocyte predictions are in **Extended Data Fig. 1C**. Amongst windows with the biggest observed cell type differences (experimental hESC versus ventricular D23; red points), CardioAkita outperforms Original Akita on 34/49 loci (69%). **D.** Exemplar genomic region from held-out test data (black point in C) with differences (black arrows) between Original Akita hESC predictions and CardioAkita ventricular D23 predictions. Differential contacts are supported by the experimental map (bottom) and include (1) lost contact at *KCNE3,* which encodes a regulatory subunit of a smooth muscle voltage-gated potassium channel, (2) gained contact at *CHRDL2* involved in bone morphogenetic protein signaling important during embryonic development.

To evaluate CardioAkita’s performance, we used held-out loci to compute two complementary metrics: mean squared error (MSE) and one minus the Pearson R correlation (1-CORR) of the predicted versus observed Hi-C maps. Along with visual inspection, these metrics indicated excellent performance across atrial and ventricular time points: mean MSE= 0.17, mean 1-CORR= 0.345. Comparing CardioAkita to Original Akita using their most similar cell types, WTC11 iPSCs and H1 human embryonic stem cells (hESCs), performance was equivalent overall (**Fig. 1B**, **Extended Data Fig. 1A**) and on a locus-by-locus basis (**Extended Data Fig. 1B**). However, CardioAkita was better able to predict atrial and ventricular chromatin interactions (**Fig. 1C**), especially in loci where the observed Hi-C data showed the greatest cell type differences (**Extended Data Fig. 1C**). For example, CardioAkita correctly predicted increased contacts for *CHRDL2* and decreased contacts for *KCNE3* at ventricular D23 versus pluripotent cells (**Fig. 1D**). This is important, because *CHRDL2* encodes a protein involved in bone morphogenetic protein signaling, which is critical for heart development^26,27^, and its downregulation is associated with hypertrophic cardiomyopathy^28^, while *KCNE3* encodes a cardiomyocyte-expressed voltage-gated potassium channel protein implicated in cardiac rhythm disorders^29,30^. Thus, training on cardiomyocyte differentiation data improved the cell-type specificity and CHD relevance of the Akita framework.

### Disruption scores are elevated in CHD

We hypothesized that CHD variants may disrupt distal enhancers or CTCF binding sites, leading to altered chromatin interactions and gene expression changes (**Fig 2A**). First, we investigated this using CardioAkita and SuPreMo to predict the impact of variants from individuals with CHD compared to unaffected controls. We used publicly available trio data and focused on *de novo* SVs (*dn*SVs), which we expect to be enriched for causal variants in a rare disease like CHD. From the CHD GENES Study of the Pediatric Cardiac Genomics Consortium^31^ (PCGC), we curated 42 high-confidence *dn*SVs (26 deletions, 16 duplications) from 34 unique probands with unknown underlying genetic pathogenesis for CHD and no history of CHD among first-degree relatives. Probands were stratified based on diagnoses into severe (conotruncal and outflow tract defects) versus non-severe CHD. As a comparative population, we used unaffected siblings from the Simons Foundation Autism Research Initiative (SFARI) Simons Simplex Collection cohort^32^ and matched each CHD *dn*SV with two control *dn*SVs based on SV type and length (**Fig. 2B**).

**Figure 2:**
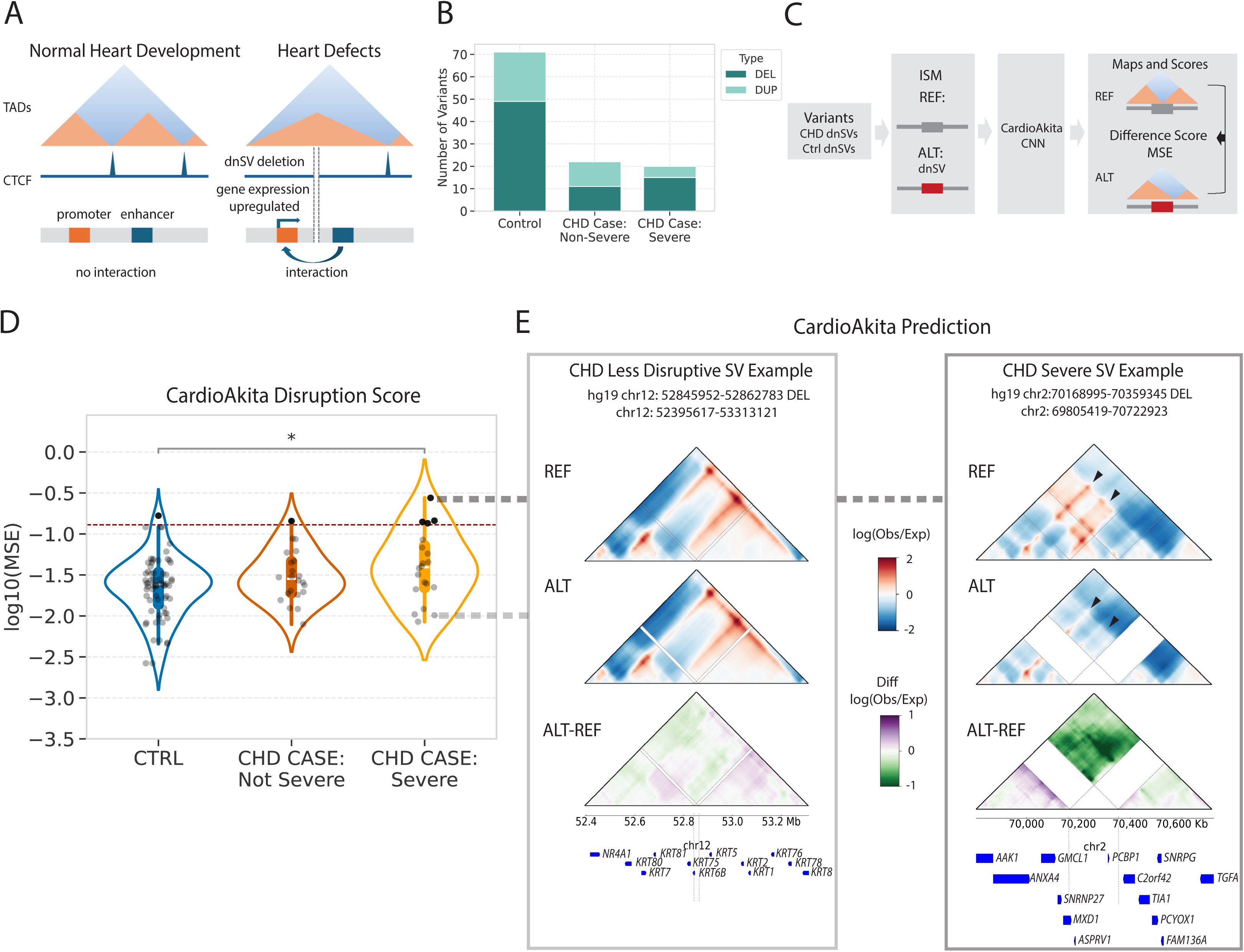
Predicted effects of *de novo* structural variants correlate with CHD severity. **A.** Proposed mechanism by which *dn*SVs disrupt chromatin boundaries (e.g., CTCF sites), leading to altered interactions and gene expression in CHD. **B.** Counts of *dn*SV types in PCGC (CHD probands grouped by severity) and SFARI SSC (unaffected controls): DEL = deletion; DUP = duplication. **C.** Schematic SuPreMo *dn*SVs scoring pipeline. Pairs of reference and alternate DNA sequences generate pairs of predicted contact maps that are compared using MSE to produce a disruption score for each CardioAkita time point. **D.** Distribution of disruption scores (log10 MSE) for *dn*SVs in **B** stratified by CHD severity. Black star: significant difference (one-sided Mann–Whitney U test); Red dotted line: 95^th^ percentile of all *dn*SVs. **E.** Predicted contact maps for two examplar *dn*SVs with low (left) and high (right) disruption scores. Dashed lines: start and end of the variant; Arrows: predicted differential genomic contacts resulting from the *dn*SV.

For each proband or control *dn*SV, we generated a pair of 1-megabase (Mb) DNA sequences, one without (reference) and one with (alternate) the variant, and predicted the corresponding pair of contact maps for each CardioAkita time point. We computed the difference (MSE and 1-CORR) between each pair of reference and alternate maps to obtain “disruption scores” for every *dn*SV at each time point (**Fig. 2C**). This cohort-wide analysis revealed greater changes to genome folding in *dn*SVs from CHD probands (**Extended Data Table 1**) compared to controls (**Extended Data Table 2**). Probands with severe CHD had *dn*SVs with the highest median disruption scores, and nearly all of the top 5% most disruptive variants were from this group **(Fig. 2D, Extended Data Fig. 2**). Visual inspection confirmed that CardioAkita’s predicted contact maps for high-scoring *dn*SVs had notable differences in chromatin loops and insulation, whereas low-scoring dnSVs induced limited changes (**Fig. 2E**). These results are the first evidence at a cohort-wide scale that CHD *dn*SVs may disrupt genome folding during cardiac development.

### Sequencing of CHD proband genomes reveals disruptive *dn*SVs

Next, we sought to evaluate CardioAkita as a tool to discover candidate causal variants in participant genomes. We selected three CHD probands newly recruited to the PCGC study based on their having no *de novo* exonic single nucleotide changes affecting known CHD-associated genes (**Extended Data Table 3**). Proband A was diagnosed with heterotaxy syndrome with polysplenia and associated cardiac defects (Ebstein’s anomaly and atrial septal defect), Proband B was diagnosed with a conotruncal defect and a ventricular septal defect, and Proband C was diagnosed with hypoplastic left heart syndrome (left-sided obstructive lesion) accompanied by mispositioning of the atrial septum. Whole-genome sequencing (WGS) of each proband and their unaffected parents enabled us to identify six *dn*SVs across the three probands: four deletions, an inversion, and a duplication (**Table 1**). We calculated disruption scores for each *dn*SV across cardiomyocyte differentiation time points (**Fig. 3**). Each proband had at least one notably disruptive *dn*SV.

**Figure 3:**
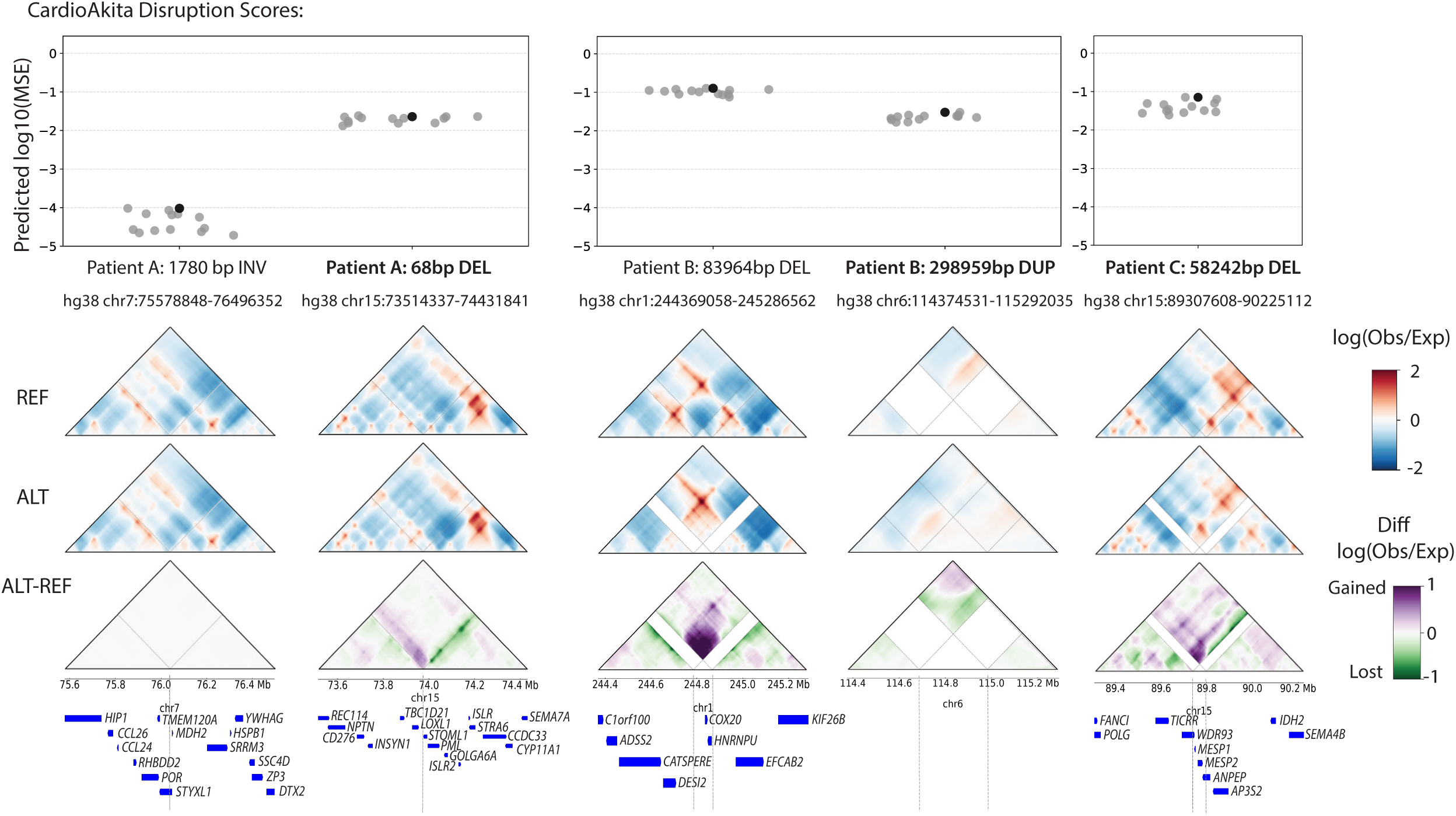
Whole-genome sequencing of CHD probands reveals *dn*SVs with high disruption scores. CardioAkita disruption scores (top) for *dn*SVs from **Table 1** (five columns; bold font: most disruptive *dn*SV per proband) with one point per timepoint (bold point: ventricular D23). Corresponding D23 contact maps (middle) show CardioAkita predictions for 1-Mb windows centered on the *dn*SV. Genes (blue bars) and *dn*SV coordinates (dashed lines) are shown below.

**Table 1:**
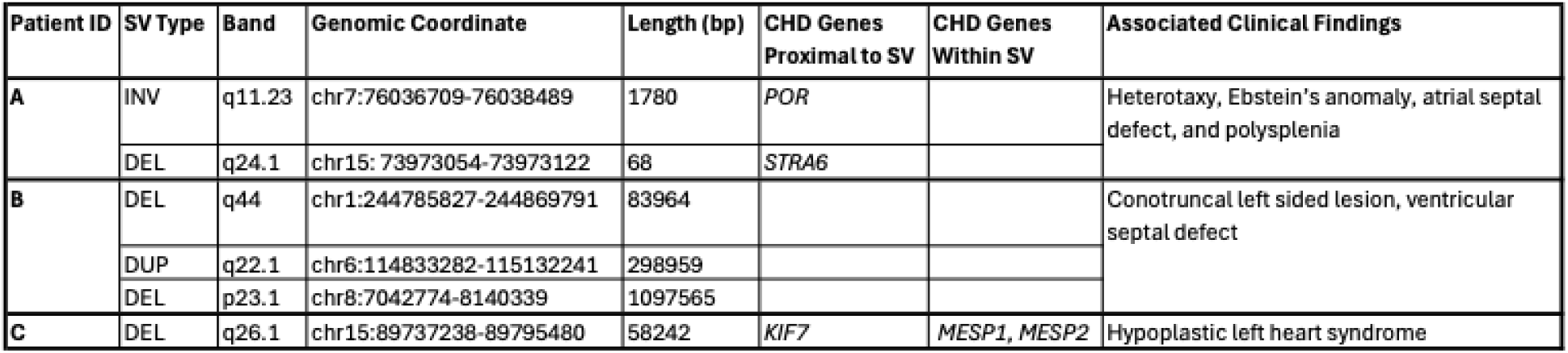
Summary of phenotypes and *dn*SVs detected in WGS of three CHD probands.

For proband A, CardioAkita predicted that a small 68-bp noncoding deletion at chromosome 15q24.1 (GRCh38 chr15:73973054-73973122) causes a loss of chromatin interactions for regions downstream of the deletion and a gain of interactions upstream of the deletion. *STRA6,* a known CHD gene that encodes a membrane receptor essential for retinoic acid signaling fundamental to heart patterning^33^, was predicted to lose interactions, whereas gained interactions were predicted for *LOXL1*, which encodes an enzyme critical for cardiac extracellular matrix homeostasis and remodeling^34^. In contrast to the 68-bp deletion, CardioAkita predicted minimal genome folding disruption for the other d*n*SV in proband A, a 1780-bp inversion at chromosome 7q11.23 proximal to the known CHD gene *POR*. These results exemplify CardioAkita’s ability to prioritize disease-relevant variants that might otherwise be considered insignificant.

For proband B, CardioAkita predicted that an 84-kb deletion at chromosome 1q44 (chr1:244785827-244869791) merges two normally insulated domains encoding *COX20* and *HNRNPU*. *COX20* is involved in the assembly of cytochrome C oxidase, and *HNRNPU* is an RNA processing gene that has been associated with developmental brain disorders, cardiomyocyte organization, and excitation-contraction coupling activities^35^. This merging of two domains was predicted to lead to ectopic interactions between the ciliary gene *EFCAB2* and the gene *DESI2,* a negative regulator of the cell cycle. Losses of insulation such as this can lead to enhancer hijacking and aberrant gene expression^36,37^. For a 300-kb noncoding duplication at chromosome 6q22.1, CardioAkita predicted loss of contact for upstream regions, but their significance is unclear because no protein coding genes were affected. Because CardioAkita makes predictions for 1-Mb windows, we could not evaluate the largest deletion in this proband at chromosome 8p23.1, a locus lacking cardiac genes.

For proband C, CardioAkita predicted that a 58-kb deletion at chromosome 15q26.1 (chr15:89737238-89795480) deletes key mesoderm and CHD-associated genes *MESP1* and *MESP2* and also merges two domains, one of which contains the CHD gene *KIF7*. As a result, we predict that this *dn*SV creates a strong ectopic contact *KIF7* and a region that includes vesicle transport gene *AP3S2*.

Thus, CardioAkita applied to WGS of three trios in which the probands lacked genetic diagnoses enabled us to prioritize several variants of unknown significance and develop testable hypotheses about how these *dn*SVs may alter chromatin and gene regulation in heart development.

### Engineered *dn*SVs from participants with CHD recapitulate CardioAkita predictions and show changes in cardiac developmental gene expression

Next, we sought to experimentally test CardioAkita’s predictions of disruptive variants in CHD probands. We selected the highest scoring *dn*SV from each genome (**Fig. 3**) to engineer into WTC11 iPSCs: proband A’s 68-bp deletion, proband B’s 84-kb deletion, and proband C’s 58-kb deletion. For the 68-bp and 58-kb deletions, we obtained homozygous lines ideal for directly comparing to our predictions, which are based on fully removing the sequence. For the 84-kb deletion, we were only able to generate a heterozygous line. All edited lines and the unedited control were differentiated into ventricular cardiomyocytes. Based on the genes neighboring each *dn*SV and CardioAkita’s predictions, we collected cells for molecular profiling at D0 (pluripotent) and D15 (cardiomyocytes) for the 68-bp deletion, D2 (cardiac mesoderm) for the 58-kb deletion, and D15 for the 84-kb deletion. We performed Capture-C sequencing to assess differential interactions around each deletion and RNA-seq to measure gene expression changes.

All three proband deletions caused changes to chromatin interactions that closely mirrored CardioAkita’s predictions, plus significant differential expression of both nearby and distal genes (**Fig. 4**, **Fig. 5**, **Extended Data Fig. 3**). Observing these effects was surprising for proband A’s *dn*SV due to its small size and intergenic location. However, it is an accessible, CTCF-bound ENCODE candidate cis-regulatory element separating two small chromatin domains. Consistent with being an insulating boundary, the deletion caused these domains to merge (**Fig. 4A**, **Extended Data Fig. 4A**). Recapitulating CardioAkita’s predictions, genes encoding three cell membrane proteins involved in developmental signaling (*ISLR2*, *ISLR*, and *STRA6*) lost interactions with regions flanking the deletion (**Fig. 4B, Extended Data Fig. 4B**). Observed differences were strongest and most concordant with predictions at D0, suggesting that the deletion plays a role early in heart development.

**Figure 4:**
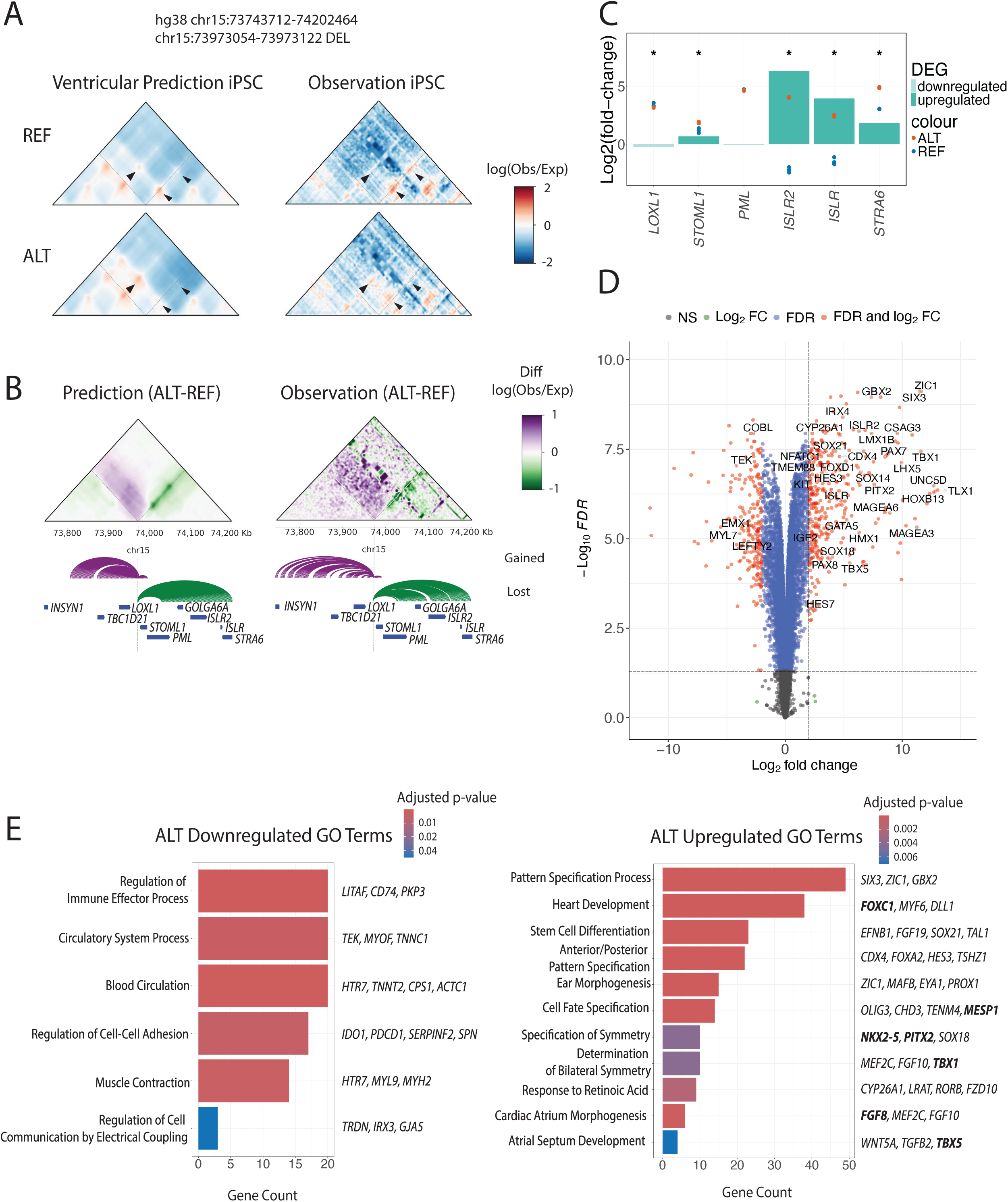
Experimental validation and gene expression effects of a 68-bp noncoding deletion. **A.** Predicted (left) and Capture-C (right) contact maps for iPSC lines unedited (top, REF) and homozygous for proband A’s 68-bp deletion (bottom, ALT). Cardiomyocyte results in **Extended Data Fig. 4**. Arrows: differential genomic contacts resulting from the *dn*SV. **B.** Differential contacts (ALT - REF). Purple: gained interactions; Green: lost interactions; Arches: differences directly flanking the deletion; Dashed lines: deletion coordinates; Blue bars: gene coordinates. **C.** Bulk RNA-seq results for genes flanking the deletion at D0. Bars: log2 fold-change in edited versus unedited lines; Black star: false discovery rate (FDR) ≤ 0.05; Blue dots: log2 counts for three unedited replicates; Orange dots: log2 counts for three edited replicates. **D.** Volcano plot of genome-wide differential expression results. Red: significant genes (FDR ≤ 0.05, log2FC > |2|); names: notable cardiac developmental genes. **E.** Gene ontology terms enriched amongst downregulated (left) and upregulated (right) genes with examples. Bold: known CHD genes; Color: FDR-adjusted p-value.

**Figure 5:**
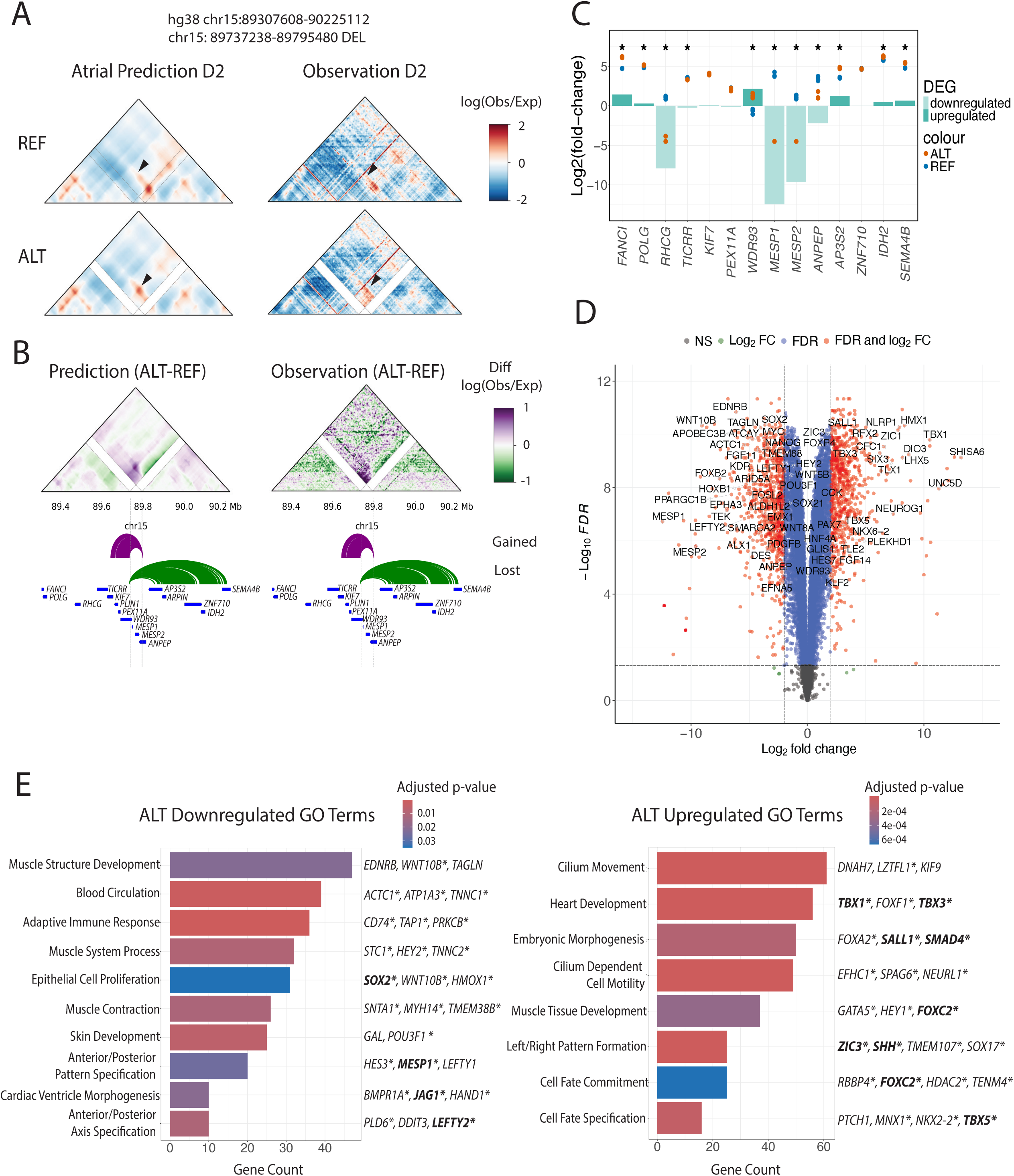
Experimental validation and gene expression effects of a 58-kbp deletion. Parallel analyses to those in Figure 4. **A.** Predicted (left) and Capture-C (right) contact maps for mesoderm (D2) cells unedited (top, REF) and homozygous for proband C’s 58-kp deletion (bottom, ALT). Results for the heterozygous deletion are in **Extended Data Fig. 5**. Arrows: differential genomic contacts resulting from the *dn*SV. **B.** Differential contacts (ALT - REF). **C.** Bulk RNA-seq results for genes flanking the deletion at D2. **D.** Volcano plot of genome-wide differential expression tests. **E.** Gene ontology terms enriched amongst downregulated (left) and upregulated (right) genes. Bold: known CHD genes; asterisks: genes also differentially expressed in heterozygous deletion lines; color: FDR-adjusted p-value.

Change in chromatin organization were associated with significant upregulation of *ISLR2*, *ISLR*, and *STRA6* at D0, up to 32-fold for *ISLR2* and decreasing with distance from the deletion (**Fig. 4C**). Transcriptome-wide, we detected upregulation of several genes involved in establishing left-right heart asymmetry (*PITX2, WNT5A, DLL1, TBX1, PITX2, FGF8, MEF2C, FGF10*) and embryonic patterning (*SIX3*, *ZIC1*, *GBX2*, *FOXC1)*, and many others (**Fig. 4D**, **Extended Data Table 4**). Several transcriptional differences are consistent with published effects of STRA6 knockdown^33^, including genes involved in mesenchyme development (*SNAI2*), retinoic acid synthesis (*ALDH1A2*), and cardiac contractility (*ACTA1*, *ACTA2*, *MYL9*). Gene Ontology (GO) analysis showed that cardiac development, atrial morphogenesis and septal development, retinoic acid response, pattern specification, cell fate commitment, and tissue morphogenesis were enriched for upregulated genes, while outflow tract septum morphogenesis and immune-related responses were enriched for downregulated genes (**Fig. 4E**, **Extended Data Table 5**).

Proband B’s 84-kb deletion was predicted to merge two domains and strengthen a chromatin loop, and this was confirmed in our Capture-C data (**Extended Data Fig. 3A**). Chromatin interactions lost upstream of the deletion (**Extended Data Fig. 3B**) included the upregulated gene *KIF26B* (**Extended Data Fig. 3C**). *KIF26B* encodes a kinesin that regulates cell migration and planar cell polarity signaling during embryonic morphogenesis^38^, and it shows correlated differential DNA methylation and expression in mouse embryonic heart development^39^. We observed widespread differential expression at D15, including downregulation of *LEFTY2*, *TFAP2A*, *SFRP2*, and *WNT* (**Extended Data Fig. 3D**, **Extended Data Table 6**). This suggests impaired early patterning and signaling precision, including left–right axis specification and neural crest competence. Concurrent reduction in *SOX2* and mesoderm-associated markers suggested diminished developmental plasticity and lineage priming, consistent with disruption of early regulatory programs rather than terminal differentiation defects. Other transcriptional changes resembled published effects of HNRNPU knockdown, including those that regulate cardiogenesis (*GARS*, *ERBB4*, *MARS*, *ASNS*) and cardiac contractility (*ACTA1*, *NPPB*)^40^. GO analysis identified downregulated pathways involved in cell fate commitment and specification, canonical WNT signaling pathway, and outflow tract septum morphogenesis, plus upregulated pathways involved in aorta and vascular development, pattern specification, extracellular matrix organization, and lipid localization. Disruption of extracellular matrix and lipid organization suggest a compensatory remodeling response to impaired developmental patterning (**Extended Data Fig. 3E**, **Extended Data Table 7**). We speculate that disruption of key lineage-defining transcription factors may stall differentiation in the deletion line, promoting enhanced cell–matrix interactions and metabolic adaptations that support membrane signaling and tissue structural integrity.

Proband C’s 58-kb deletion had the most obvious connection to CHD, because it overlaps *MESP1* and *MESP2,* important regulators heart development^41–43^. However, CardioAkita’s prediction of broad changes to chromatin organization in the locus suggested that the deletion’s effects might be more complex than loss of the MESP transcription factors. We focused on D2, which corresponds to the peak expression of *MESP1,* corresponding to early mesoderm specification and cardiac lineage commitment^41,44^. Capture-C results recapitulated the CardioAkita prediction of a shift in the location of a strong chromatin loop anchor (**Fig. 5A**), involving both gained and lost interactions (**Fig. 5B**).

Upstream of the deletion, we observed increased gene expression for *AP3S2*, *IDH2*, and *SEMA4B,* which correlated with their loss of chromatin contacts (**Fig. 5C**). *IDH2* and *SEMA4B* are involved in cardiac metabolism, maturation, and neural crest positioning and signaling^45,46^. We also observed gained contacts and downregulation of *ANPEP,* a gene involved in endothelial and vascular development^47^. Downstream of the deletion, we observed gained contacts, accompanied by decreased expression of *RHCG* and *TICRR* and increased expression of *FANCI*, *POLG*, and *WDR93* (**Fig. 5B, 5C**). Beyond these local expression changes, we observed widespread transcriptional changes involving known cardiac developmental genes (**Fig. 5D**, **Extended Data Table 8**). GO analysis further highlighted disruption of developmental programs, with upregulated pathways related to cilium movement and organization, cardiac morphogenesis, and pattern specification, and downregulated gene pathways enriched for anterior/posterior axis specification and inflammatory response processes (**Fig. 5E**, **Extended Data Table 9**). We found that this deletion resulted in many transcriptional changes consistent with MESP1 knockdown including emerging cardiac mesodermal genes (*EOMES*, *LHX1*, *MIXL1*), early gastrulation (*UPP1*, *GSC*), and cardiac structural genes (*MYL7*, *ACTA2*)^42^. These results suggest that loss of *MESP1/MESP2* and/or the differential expression of flanking genes may have perturbed regulatory networks governing cardiac fate decisions. Given the role of *MESP1* in cardiovascular progenitor specification^48^, it is difficult to determine if the differential chromatin interactions and expression of flanking genes also contribute to the global expression differences in the deletion line. Genetically or epigenetically isolating these effects is a challenging but important direction for future work.

As an initial step in this direction, we used CRISPR/Cas9 to genetically engineer heterozygous WTC11 iPSC lines carrying the 58-kb deletion, differentiated them to the mesoderm stage, and performed RNA-seq. We expected to observe attenuated effects compared to the homozygous deletion, which would be important since heterozygosity reflects the allelic state in proband C. Similar to the homozygous deletion, we observed a local upregulation of *SEMA4B* and *IDH2* (**Extended Data Fig. 5A**), plus global transcriptional changes (**Extended Data Fig. 5B**, **Extended Data Table 10**) that included upregulation of critical cardiac transcription factors (*TBX1*, *TBX5*, *FOXF1*, *GATA5*) and downregulation of early developmental genes (*HAND1*, *LEFTY2*, *FOXB2)*. Differentially expressed genes were similarly enriched for roles in cell fate specification, pattern specification, and tissue morphogenesis functions (**Extended Data Fig. 5C-D**, **Extended Data Table 11**). These results demonstrate that the 58-kb deletion is haploinsufficient, as are many developmental disorder genes^49^, and that it plays an early and precise role in regulating cardiac progenitor specification that may be regulated in part by 3D chromatin organization.

While further work is necessary to characterize the *dn*SVs in this study and evaluate if they play causal roles in CHD, our results clearly demonstrate that CardioAkita can accurately predict the effects of variants from people with CHD on 3D genome organization and prioritize variants with widespread effects on disease-relevant gene expression.

## Discussion

As a machine learning model, Akita has significantly furthered our understanding of how sequence disruptions resulting from SVs can affect 3D genome organization across several different biological contexts including evolution, pediatric brain tumors, and autism^20,23,50^. For CHD, while copy number variants are known to contribute to as many as 10-15% of CHD cases, the underlying mechanisms through which they cause CHD are largely unknown, especially for intergenic variants^31,51^. To accelerate discovery of novel CHD variants and mechanisms, we created CardioAkita, improving our ability to predict chromatin interactions that are specific to differentiating atrial and ventricular cardiomyocytes. Leveraging this model, we established a pipeline for screening WGS data and generating testable hypotheses regarding variants that disrupt chromatin and gene regulation. This led to experiments on variants that would likely be considered insignificant if identified in clinical sequencing yet had profound effects on cardiac gene expression programs, including causing significant differential expression of key regulators of cell fate, differentiation, and heart development. In addition, our analysis of dozens of *dn*SVs revealed a positive association between CardioAkita disruption scores and CHD severity. These findings provide the first evidence that disrupted 3D genome organization may play a mechanistic role in CHD.

This investigation has several limitations. First, our cohort-level analysis required using unrelated individuals from SFARI SCC, because PCGC did not sequence unaffected siblings. We felt that SCC siblings were a conservative choice of unaffected controls (as opposed to using 1000 Genomes unaffected trios, for example), because having a sibling with autism means that these individuals are more likely than the general population to possess damaging variants^23^. Even though ancestry has only a weak association with *de novo* mutation rate that may be environmentally mediated^52^, another factor in using SCC was the distribution of genetic ancestries being similar to PCGC. We also matched *dn*SVs for type and length. Nonetheless, technical biases between SCC and PCGC could affect our results. Second, training Akita-style models requires very high-resolution Hi-C. This limited the developmental stages and cardiac cell types we could model. The differentiated cardiomyocytes we utilized are relatively immature compared to adult cardiomyocytes, so genome organizational differences that are uniquely present in immature cardiomyocytes will be weighed more heavily in our analysis. This potentially limits the specificity of CardioAkita for other types of cardiovascular diseases, including those with an adult onset. However, there is significant evidence that among cardiovascular diseases affecting adults, such as heart failure, cardiomyocytes revert to a fetal phenotype^53,54^. This limitation is also attenuated because studies have shown that over 60% of SVs with gene expression effects impact several tissue types^55^. Nonetheless, training with additional Hi-C data would make CardioAkita more generalizable. Finally, our experiments primarily used homozygous deletions, while the *dn*SVs were heterozygous in probands. Hence, further work is needed to characterize heterozygous variant effects.

Beyond the specific datasets and model presented here, our findings highlight the profound implications off emerging machine-learning approaches on improving the quality of genetic diagnosis and on our ability to quickly ascertain the relative pathogenicity and biological consequences of variants. Being able to predict the molecular effects of gene regulatory variants with cell type specificity and in the context of different genetic backgrounds will ultimately improve our ability to interpret the potential pathogenicity of clinical WGS data. Computational variant effect prediction is useful, because it is low cost, efficient, and capable of generating testable hypotheses about molecular mechanisms, enabling the vast number of potential regulatory variants to be prioritized for follow up. This is particularly important in contexts where the feasibility of experimental characterization is limited.

## Online Methods

### CardioAkita training and evaluation

Data for training CardioAkita was generated across discrete differentiation time points from inducible pluripotent stem cells into atrial or ventricular cardiomyocytes as previously described^24^. The ventricular time points were day (D) 0 (iPSC), D2 (mesoderm), D4 (cardiac mesoderm), D6 (cardiac precursors), D11 (early cardiomyocytes), and D23 (later cardiomyocytes). The atrial time points were D2 (mesoderm), D4 (cardiac mesoderm), D6 (cardiac precursors), D11 (early cardiomyocytes), D20 (later cardiomyocytes), D45 (later cardiomyocytes), and the same D0 (iPSC) timepoint as the ventricular time series.

Hi-C sequencing reads were mapped to GRCh38 (also known as hg38) and processed into bins 2048 base pairs (bp) long, followed by genome-wide iterative correction (ICE) normalization, adaptive coarse-graining, observed over expected normalization, log transformation, clipping to (−2,2), linear interpolation, and convolution with a 2D Gaussian filter for smoothing. These data were segmented into windows ∼ 1 megabase (Mb) long (2^20^ bp) and paired with the underlying DNA sequences.

The CardioAkita model has the same loss function as the published Orginal Akita model^21^, and the convolutional neural network architecture is similar except that the output head makes predictions for each atrial and ventricular time point. We compared performance of an architecture that jointly predicts all atrial and ventricular time points versus one trained separately on each differentiation, and we found predictions to be most accurate for the separate approach. From a user perspective, the CardioAktia prediction framework takes a ∼ 1 Mb sequence as input and predicts Hi-C for all twelve time points. This is accomplished using an atrial-trained and a ventricular-trained model.

To train CardioAkita, we split genome-wide windows split into training (n=7008), validation (n=419) and test (n=413) sets. Training windows were used to estimate model parameters, while validation windows were used during training to select hyperparameters (e.g., early stopping). Test windows were used to evaluate model performance on parts of the genome never seen during training.

To evaluate CardioAkita’s performance, we used two complementary metrics: mean squared error (MSE) and Pearson’s correlation (1-CORR) from the python SciPy library. These metrics quantify how much the predicted Hi-C interaction map differs from the corresponding observed map for a given window. MSE tends to capture deviations in the strength of chromatin interactions, whereas 1-CORR is more sensitive to differences in the pattern of chromatin interactions. The distribution of these metrics across held-out test windows provides an assessment of the model’s ability to make accurate and generalizable predictions. To focus performance evaluations on windows with cell type differences, we used MSE to select a subset of windows with the largest differences between pairs of observed chromatin interactions maps (MSE > 0.25, which is one standard deviation above the mean predicted versus observed performance for both Original Akita and CardioAkita).

Since Original Akita does not make predictions for cardiomyocytes but does make predictions for human embryonic stem cells (H1 hESCs), we performed head-to-head comparisons using Orginal Akita’s hESC predictions and CardioAkita’s iPSC predictions, the most closely matched cell types. To test if CardioAkita improved upon Orginal Akita’s ability to make accurate predictions for cardiomyocyte time points, we also used the observed atrial D45 and ventricular D23 Hi-C data for held-out test windows to evaluate predictions from CardioAkita for these time points against predictions from Orginal Akita’s different cell type outputs: hESC, human foreskin fibroblast (HFF) cells, B-lymphoblastoid (GM12878) cells.

### Structural variant identification for wide-scale profiling and variant exclusions

CHD trios (n=538 families with a proband and two unaffected parents) were previously recruited into the CHD GENES Study of the Pediatric Cardiac Genomics Consortium (PCGC). We classified the following CHD phenotypes as “severe”: left ventricular outflow tract obstruction (hypoplastic left heart syndrome); left ventricular outflow tract obstruction (atrial septal defect/hypoplastic left heart syndrome); conotruncal defects (tricuspid atresia/dextro-transposition of the great arteries); conotruncal defects (tetralogy of Fallot/absent pulmonary valve syndrome); conotruncal defects (tetralogy of Fallot/pulmonary atresia); conotruncal defects (tetralogy of Fallot/dextro-transposition of the great arteries); conotruncal defects (dextro-transposition of the great arteries/ventricular septal defect); and conotruncal defects (dextro-transposition of the great arteries). CHD probands without any of these phenotypes were classified as “not severe”.

In the CHD GENES Study, individuals in each trio were genotyped using genome-wide dense single nucleotide polymorphism arrays and whole exome sequencing, and *dn*SVs were called using PennCNV. For this study, we conservatively chose to only analyzed the sixty-three proband *dnS*Vs that were validated by the PCGC using digital droplet polymerase chain reaction experiments^56–58^.

Since PCGC did not genotype siblings, we evaluated different sources of control *dn*SVs. SVs from unrelated individuals filtered to singletons or extremely low frequency in population cohorts like gnomAD or 1000 Genomes would provide a large number of unaffected individuals that could be matched to PCGC probands based on genetic ancestry and biological sex, but without parental sequencing it is difficult to know if these rare variants are comparable to true *dn*SVs. So, we considered using trios from 1000 Genomes^59^ or simplex families from the Simons Foundation Autism Research Institute (SFARI) SCC cohort^32^, which include unaffected siblings. After comparing dnSV counts, length distributions, and genomic distributions, we chose SFARI siblings as the most conservative option. These individuals have siblings with autism, and hence are expected to carry a higher burden of damaging variants than the general population^23^. The *dn*SVs were previously called for SCC unaffected siblings (n=348) from short-read whole genome sequencing (WGS) data using the GATK-SV pipeline^60^.

To evaluate *dn*SVs using CardioAkita, we needed to focus on variants smaller than the ∼1-Mb prediction window. Hence, *dn*SVs greater than 700 kilobases (kb) in length were filtered out. We also required that each *dn*SV had a known alternate allele sequence and was present in the GRCh38 assembly. For each of the remaining 42 CHD *dn*SVs, we selected up to two control *dn*SVs that were matched on length (within 10%) and structural variant type. This generated 67 control *dn*SVs.

### Whole-genome sequencing

Three trios from the PCGC cohort were selected for genome-wide *dn*SV discovery using WGS. They were prioritized based on the proband having a severe diagnosis and no predicted damaging amino acid changes in known CHD genes. WGS was completed on trios (participant with CHD and parents), as previously described^61^. In brief, genomic DNA samples from blood or saliva were prepared for sequencing using a polymerase chain reaction-free library preparation or SK2-IES library preparation. All samples were sequenced on an Illumina Hi-Seq X Ten with 150-bp paired-end reads to a median depth >30x per individual.

The *dn*SVs from these selected trios were called by pooling three pipelines from PCGC members at Harvard Medical School and the University of Utah. At Harvard, multiple tools and types of SV evidence were used to call *dn*SVs from trio genome sequencing. *De novo* deletions and duplications were called from alternations in phased B-allele frequency and allele depth using MoChA (https://github.com/freeseek/mocha). WGS VCF files were phased with Eagle (v2.4.1) using the 1000 Genomes reference panel, incorporating pedigree information to improve haplotype phasing, followed by variant calling with the MoChA plugin for Bcftools (v1.9). SVs were also called using genome-wide local assembly with SvABA (FH Version: 134)^62^. SvABA was run for each trio using the proband BAM files as ‘case-bam’ and parental BAM files as ‘control-bam’. Regions of the genome with poor mappability (https://www.encodeproject.org/annotations/ENCSR636HFF/) were excluded from SV calling.

Primary evidence supporting SV calls was visualized for each trio in IGV to evaluate the quality of evidence supporting the call and confirm the *de novo* status of each variant. At the University of Utah, the reference-free RUFUS algorithm (https://github.com/jandrewfarrell/RUFUS/) was applied to the WGS data for *de novo* variant calling, and results were filtered to retain SVs >50 bp in length.

### Variant scoring

For variant scoring with CardioAkita, we used a computational pipeline called Sequence Mutator for Predictive Models (SuPreMo) to streamline *in silico* perturbations^22^. For each structural variant, SuPreMo generates a ∼1-Mb reference sequence centered on the variant’s coordinate and a corresponding sequence with the variant’s alternate allele incorporated at the appropriate position. These sequences are automatically fed into CardioAkita, which produces predicted chromatin contact maps for both sequences at each time point. SuPreMo pads and masks the resulting maps to align the 2048-bp bins in each pair of maps. Regions of the maps flanking the *dn*SV are then compared to each other (reference prediction versus alternate allele prediction) using MSE and 1-CORR. To ensure the robustness of these disruption scores, we implemented augmentation by additionally predicting and scoring maps after modifying the input sequence by shifting 1 bp right, shifting 1 bp left, and reverse complementing. For each *dn*SV, we reported the average across the four scores (no augmentation and three augmentations).

### Cell line generation

WTC11 iPSCs (Coriell #GM25256), a gift from Dr. Bruce Conklin, were used in this study. The CRISPR (clustered regularly-interspaced short palindromic repeats)/Cas system was used by Gladstone Institute’s Stem Cell Core (San Francisco, CA) and Sampled (Piscataway, New Jersey) to generate edited cell lines for three CardioAkita prioritized CHD *dn*SVs: a 68-bp homozygous deletion (chr15: 73973054-73973122), a heterozygous and a homozygous 58-kb deletion (chr15: 89737238-89795480), and a heterozygous 84-kb deletion line (chr1: 244785827-244869791). iPSC cell lines were electroporated with Cas9 protein, synthetic guide RNA, and a donor oligo containing the desired repair. Cells were plated as single cells in 96 well plates for screening. Successfully edited cells were identified using Sanger sequencing and subsequently expanded. Additional quality control measures such as mycoplasma testing, evaluating for unintended off-target modifications, and monitoring for cell line homogeneity were performed.

### Maintenance of iPS cells and differentiation

iPSC protocols were approved by UCSF’s Human Gamete, Embryo, and Stem Cell Research Committee and the Institutional Review Board. Differentiation for WTC11 iPSCs used for atrial and ventricular time course HiC 3.0 experiments were performed as previously described^24^. Human iPSCs for all other experiments were plated on growth factor-reduced Matrigel basement membrane (Corning #356231) and grown in mTESR-1 (StemCell Technologies #85850) or mTESR-Plus (StemCell Technologies #100-0276). ReLESR (StemCell Technologies #100-0483) was used for routine passaging and Accutase (StemCell Technologies #07920) was used for iPSC dissociation. For differentiation, cells were grown for three days until reaching 90% confluency and induced using Stemdiff Cardiomyocyte Ventricular Kit (StemCell Technologies #05010) according to manufacturer’s instructions.

### Capture-C library preparation and sequencing

We processed two biological replicates for Capture-C library preparation for each *dn*SV engineered line and the corresponding isogenic control lines (transfected with non-targeting guide RNA at D0, D2 and D15 respectively). For each biological replicate, cells were grown in a 10-cm dish and were harvested using 0.25% Trypsin-EDTA for 5 minutes (min), singularized by repeated resuspension using P1000 pipette, followed by quenching in cell culture media. Ten million cells were crosslinked with 37% formaldehyde to bring the final concentration to 2% formaldehyde and incubated at room temperature for 10 min. Samples were subsequently quenched with 2.5M glycine to a final concentration of 200mM and incubated at room temperature for 5 min. Cells were pelleted at 500 x g for 5 min at room temperature. Supernatant was carefully removed and cell pellets were snap frozen in liquid nitrogen and stored at -80°C. Capture-C data was generated by Arima Genomics according to their standard protocol described in the Arima-HiC+ (A510008), Library Prep Module (A303010), and the Capture Module kits (A302010). SureDesign custom probes were ordered from Agilent (Santa Clara, CA). Target enrichment was performed using a custom panel targeting the following ∼2-Mb *dn*SV-centered regions (GRCh38): chr15: 88766359-90766359, chr1: 243827809-245827809, and chr15:72973088-74973088. Libraries were sequenced on an Illumina NovaSeq 6000 instrument using S4 flow cells.

### Capture-C data analysis

The 4DN pipeline (https://data.4dnucleome.org/resources/data-analysis/hi_c-processing-pipeline) was used to process Capture-C sequencing libraries. Reads were mapped to GRCh38 using bwa v0.7.17^63^. Valid Capture-C alignments were filtered using pairtools v0.3.0 and compiled into a pairs file^64^. After confirming high replicate concordance, the biological replicates of each cell line were merged using the 4DN run-merge-pairs script (https://github.com/4dn-dcic/docker-4dn-hic/blob/master/scripts/). The pairs files were converted to cool files using Cooler v0.8.11^65^. To visualize Capture-C contact frequency maps, cool files were preprocessed at a bin resolution of 2048 bp in the same way as the datasets used to train the CardioAkita model^21,66^. Specifically, Capture-C data was normalized with genome-wide ICE, matrices were smoothed using adaptive coarse graining from cooltools v0.7.0^67^, and normalized for distance-dependent contact decay^25^. The values were log scaled and truncated to the range (−2,2). Matrices were linearly interpolated to fill missing values using cooltools v0.7.0. Maps were smoothed using astropy v6.1.3 convolution with a 2D Gaussian filter.

### Bulk RNA-sequencing generation

We profiled gene expression for the homozygous 68-bp deletion on chr15 at D0 (n = 2), the homozygous 58-kb deletion on chr15 at D2 (n = 4), the heterozygous 58-kb deletion on chr15 at D2 (n = 3), and the heterozygous 84-kb deletion at D15 (n=2). For each line, we used the same number of replicates from corresponding control lines transfected with a non-targeting guide RNA. RNA was isolated using Qiagen RNeasy Kit (Cat. No. 74104) according to manufacturer’s instructions. RNA samples were processed using the Illumina Stranded mRNA Prep Ligation Kit (20040532) according to manufacturer’s instructions. For each sample, 250ng of total RNA (quantified on the Invitrogen Qubit 2.0 Fluorometer using the Qubit RNA HS Assay Kit (Q10210)) was subjected to purification using oligo(dT) magnetic beads to capture the polyadenylated mRNA molecules. Following purification, RNA was primed with random hexamers for first-strand cDNA synthesis, and then second-strand cDNA synthesis. During second-strand cDNA synthesis, deoxyuridine triphosphate (dUTP) was incorporated in place of deoxythymidine triphosphate (dTTP) to achieve strand specificity in a subsequent amplification step. Next, adenine (A) nucleotide was added to the 3’ ends of the blunt fragments to prevent ends from ligating to each other and to provide a complementary overhang to the thymine (T) nucleotide on the 3’ end of the adapter. During adapter ligation and amplification, indexes and adapters were added to both ends of the fragments, resulting in 10-bp, dual-indexed libraries, ready for cluster generation and sequencing. The second-strand was quenched during amplification due to the incorporation of dUTP during second-strand cDNA synthesis, allowing for only the antisense strand to be sequenced in Read 1. Twelve cycles of amplification were performed. Each library was run on the Agilent Bioanalyzer, using the Agilent High Sensitivity DNA Kit (5067-4626), to assess the size distribution of the libraries. They were quantified on the Invitrogen Qubit 2.0 Fluorometer using the Qubit 1X dsDNA HS Assay Kit (Q33230). Each library was normalized, then pooled equimolarly. Pooled libraries were submitted to the University of California San Francisco Center for Advanced Technology (UCSF CAT) for one lane of sequencing on the Illumina NovaSeqX 10B 100 flow cell, with run parameter 50x10x10x50bp.

### Bulk RNA-sequencing analysis

RNA reads for cell lines carrying each proband deletion and corresponding unedited control lines were processed using the nf-core/rnaseq Nextflow pipeline^68^, which performs transcript quantification using Salmon^69^. Using the *filterByExpr* function implemented in edgeR package^70^, we filtered out long noncoding RNA genes and genes with low raw counts in 30% or more of samples using a threshold empirically selected based on plots of variance versus mean expression (min.count=70 for the 68-bp deletion, 200 for the 84-kb deletion, and 60 for the 58-kb deletion). The number of retained, expressed genes was 13,033 for the 68-bp homozygous deletion, 12,716 for the 84-kb heterozygous deletion, 12,245 for the 58-kb homozygous deletion, and 12,185 for the 58-kb heterozygous deletion. After adding a pseudocount of 0.01, we estimated logCPM of each expressed gene in each sample. To visually assess if deletion lines and control lines clustered separately from each other, we performed Principal Component Analysis (PCA) on the logCPM values using the *prcomp* R package. All comparisons showed differentiation between the genotypes (data not shown). Next, reads were normalized using the *calcNormFactors* function, the dispersion was estimated using *estimateDisp* function, and the normalized reads were modeled using *glmQLFit* function implemented in edgeR package. Differential expression was defined as log fold-change between genotypes (logFC) > |2| and false discovery rate (FDR) adjusted p-value ≤ 0.05^71^. Volcano plots generated with the EnhancedVolcano R package^72^ were used to visualize differentially expressed genes.

Overrepresentation analysis was performed separately for sets of upregulated (logFC > 2) and downregulated (logFC < -2) genes against the Gene ontology (GO) database^73^ using the *enrichGO* function implemented in the clusterProfiler R package^74^. For comparing deletions to controls, the background set was all expressed genes in that comparison. Differentially expressed genes that were shared between the 58-kb heterozygous and homozygous lines were compared against a background of all genes expressed in both heterozygous and homozygous cell lines.

## Acknowledgements

We thank Mylinh Bernardi and members of the Gladstone Stem Cell Core, Gladstone Flow Cytometry Core and the UCSF Center for Advanced Technologies for their expert assistance. This work was supported by grants from the National Institutes of Health (NIH 4D Nucleome Project NHLBI U01HL157989 to B.G.B. and K.S.P.; NHLBI Bench to Bassinet Program UM1HL098179 to K.S.P. and B.G.B); Additional Ventures Innovation Fund to K.S.P and B.G.B.; Keck Foundation to K.S.P.; and Milken Institute/Biswas Family Foundation to K.S.P. Other support was provided by the following gifts and awards: NHLBI TOPMed Fellowship to J.L.; NHLBI award F31HL156439 to M.P.; Additional Ventures Catalyst to Independence Award to Z.L.G.; Chan Zuckerberg Biohub to K.S.P.; Whittier Foundation to K.S.P.; Younger Family Fund to B.G.B.; Rodenberry Foundation to B.G.B.; and California Institute for Regenerative Medicine to J.W. Sequencing was performed at the UCSF CAT, supported by UCSF PBBR, RRP IMIA, and NIH 1S10OD028511-01 grants.

## Author Contributions Statement

M.P. and K.S.P. conceived the study; J.L. developed CardioAkita; J.L.; S.U.M. and the Pediatric Cardiac Genomics Consortium performed whole-genome sequencing; M.P. and G.F. made predictions for patient variants; J.W., R.K. and RUCDR Stem Cell Services performed genome editing; J.W., Z.L.G. and K.H. performed stem cell differentiations and Capture-C and RNA-seq library preparation; Arima Genomics performed Capture-C probe design and sequencing; J.L., S.K. and M.P. analyzed Capture-C data; J.L. and M.T. analyzed RNA-seq data; J.L. performed Gene Ontology analyses; J.L., J.W., B.G.B. and K.S.P. interpreted results and wrote the manuscript; all authors reviewed and edited the manuscript.

## Extended Data Figure Captions

**Extended Data Figure 1: CardioAkita predicts 3D genome folding in differentiated cardiomyocytes from DNA sequence. A.** Reexhibited bar plot from Fig. 1 with iPSC predictions (atrial model). Bar plot representing the mean performance across 413 held-out genomic windows with error bars denoting +/- 1 standard deviation for Original Akita hESC predictions and CardioAkita iPSC predictions (ventricular and atrial models). Performance measured by mean squared error (MSE) and 1 minus Pearson R (1-CORR) between observed and predicted chromatin contact maps. **B.** Comparison of model performance between CardioAkita iPSC versus Original Akita hESC for each of the 413 held-out genomic windows from the test set. Each window is represented as a blue dot. Left column: CardioAkita iPSC performance (atrial model) plotted against Original Akita hESC performance. Right column: CardioAkita iPSC performance (ventricular model) plotted against Original Akita hESC performance. **C.** Performance across 413 genomic windows from the held-out test set, where each genomic window is represented as a dot. Top Section (atrial): MSE (above) and 1-CORR (below) between observed atrial D45 Hi-C contact maps and CardioAkita’s atrial D45 predictions (vertical axes) plotted against MSE and 1-CORR between observed atrial D45 Hi-C contact maps and Original Akita’s predictions for H1 hESC (column 1), HFF (column 2), and GM12878 (column 3) cells. Bottom Section (ventricular): MSE (above) and 1-CORR (below) between observed ventricular D23 Hi-C contact maps and CardioAkita’s ventricular D23 predictions (vertical axes) plotted against MSE and 1-CORR between observed ventricular D23 Hi-C contact maps and Original Akita’s predictions for H1 hESC (column 1), HFF (column 2), and GM12878 (column 3) cells. Red dots represent genomic windows with the largest cell type differences in the observed data, where observed H1 hESC, HFF, and GM12878 Hi-C contact maps differed the most from observed ventricular D23 Hi-C contact maps (MSE>0.25).

**Extended Data Figure 2**: ***De novo* structural variants associated with congenital heart disease demonstrate greater disruption to 3D genome folding.** Distribution of disruption scores (log10 MSE) for each *dn*SV from SFARI controls and CHD cases stratified by severity. Disruption scores quantified using SuPreMo and predicted by CardioAkita. Panels correspond to atrial (top) and ventricular (bottom) time points. Within each panel, each dot represents a *dn*SV. Black star indicates a statistically significant difference between the disruption scores for *dn*SVs in severe CHD cases compared to controls, calculated using a one-sided Mann–Whitney U test of the null hypothesis of equal distributions versus the alternative of higher scores in sever CHD cases.

**Extended Data Figure 3: Experimental Capture-C map validates predicted genomic contact map from CHD proband C’s chromosome 1q44 84-kb heterozygous deletion. A.** Left: Predicted ventricular early cardiomyocyte (D11) contact map predictions for the reference genome (REF) and the reference genome modified to have proband C’s 84-kb heterozygous deletion (ALT). Right: Observed early cardiomyocyte (D15) Capture-C contact map for cells without (REF) and with (ALT) the heterozygous *dn*SV. Differential genomic contacts between REF and ALT as a result of the *dn*SV are highlighted by black arrows. **B.** Difference maps (ALT - REF) for predicted and observed contact maps. For regions flanking the deletion, gained (purple arches) and lost (green arches) genomic contacts are represented below. Vertical dashed lines represent the start and end of the variant. **C.** Bulk RNA-seq results at D15 showing the log2 fold-change of differentially expressed genes within the 1-Mb window centered on the 84-kb heterozygous deletion. Blue dots represent the log2 fold-change from two REF replicates, and orange dots represent the log2 fold-change from two ALT replicates. Black stars represent statistically significant differentially expressed genes. **D.** Transcriptome-wide D15 bulk RNA-seq results shown as a volcano plot comparing ALT and REF lines. Significant (FDR ≤ 0.05 and log2FC > 2) upregulated and downregulated genes are shown in red with notable cardiac developmental genes named. **E.** Gene ontology analysis of differentially expressed genes showing the most enriched GO terms, the number of upregulated or downregulated genes annotated to each term (bar), FDR-adjusted p-values (color), and names of exemplar genes.

**Extended Data Figure 4: Experimental Capture-C map validates predicted genomic contact map from CHD proband A’s chromosome 15q24.1 68-bp homozygous deletion. A.** Left: Predicted ventricular early cardiomyocyte (D11) contact map prediction for the reference genome (REF) and the reference genome modified to have proband A’s 68-bp homozygous deletion (ALT). Right: Observed early cardiomyocyte (D15) Capture-C contact map for cells without (REF) and with (ALT) the *dn*SV. Differential genomic contacts between REF and ALT as a result of the *dn*SV are highlighted by black arrows. **B.** Difference maps (ALT - REF) for predicted and observed contact maps. For regions flanking the deletion, gained (purple arches) and lost (green arches) genomic contacts are represented below. Vertical dashed lines represent the start and end of the variant. This figure parallels Fig. 4, but shows results for D11 rather than D0.

**Extended Data Figure 5: Differentially expressed genes shared between heterozygous and homozygous lines for proband B’s 58-kb deletion. A.** Bulk RNA-seq results at D2 showing the log2 fold-change of expressed genes within the 1-Mb window centered on proband B’s 58-kbp heterozygous deletion. Blue dots represent the log2 fold-change from three REF replicates, and orange dots represent the log2 fold-change from three ALT replicates. Black star represents statistically significant differentially expressed genes. **B.** Transcriptome-wide D2 bulk RNA-seq results shown as a volcano plot comparing heterozygous ALT and REF lines. Significant (FDR ≤ 0.05 and log2FC > 2) upregulated and downregulated genes are shown in red with notable cardiac developmental genes named. Panels A and B parallel Fig 5, but show results for an engineered heterozygous deletion, rather than a homozygous deletion. **C.** Gene ontology analysis of differentially expressed upregulated (top) and downregulated (bottom) genes that are shared between the heterozygous and homozygous lines. Plots show the most enriched GO terms, the number of upregulated or downregulated genes annotated to each term (bar), FDR-adjusted p-value (color), and names of exemplar genes.

## Extended Data Tables

**Extended Data Table 1: CardioAkita disruption scores for 42 dnSVs from the PCGC CHD GENES Study (affected individuals).**

**Extended Data Table 2: CardioAkita disruption scores for 67 dnSVs from the SFARI SCC Study (unaffected individuals).**

**Extended Data Table 3: CardioAkita disruption scores for 5 dnSVs ascertained using whole-genome sequencing of three individuals affected by CHD.**

**Extended Data Table 4: Bulk RNA-seq differential expression results for homozygous 68-bp deletion versus the isogenic unedited control at D0.**

**Extended Data Table 5: Gene Ontology over-representation analysis results for differentially expressed genes from Extended Data Table 4 (68-bp homozygous deletion at D0).**

**Extended Data Table 6: Bulk RNA-seq differential expression results for 84-kb heterozygous deletion versus the isogenic unedited control at D15.**

**Extended Data Table 7: Gene Ontology over-representation analysis results for differentially expressed genes from Extended Data Table 6 (84-kb heterozygous deletion at D15).**

**Extended Data Table 8: Bulk RNA-seq differential expression results for 58-kb homozygous deletion versus the isogenic unedited control at D2.**

**Extended Data Table 9: Gene Ontology over-representation analysis results for differentially expressed genes from Extended Data Table 8 (58-kb homozygous deletion at D2).**

**Extended Data Table 10: Bulk RNA-seq differential expression results for 58-kb heterozygous deletion versus the isogenic unedited control at D2.**

**Extended Data Table 11: Gene Ontology over-representation analysis results for differentially expressed genes from Extended Data Table 10 (58-kb heterozygous deletion at D2).**

