## Supplementary figures and images for "Structural variants in human congenital heart disease disrupt distal genomic regulatory contacts of developmental genes"

### Extended Data Figure 1

Supplementary Figure 1

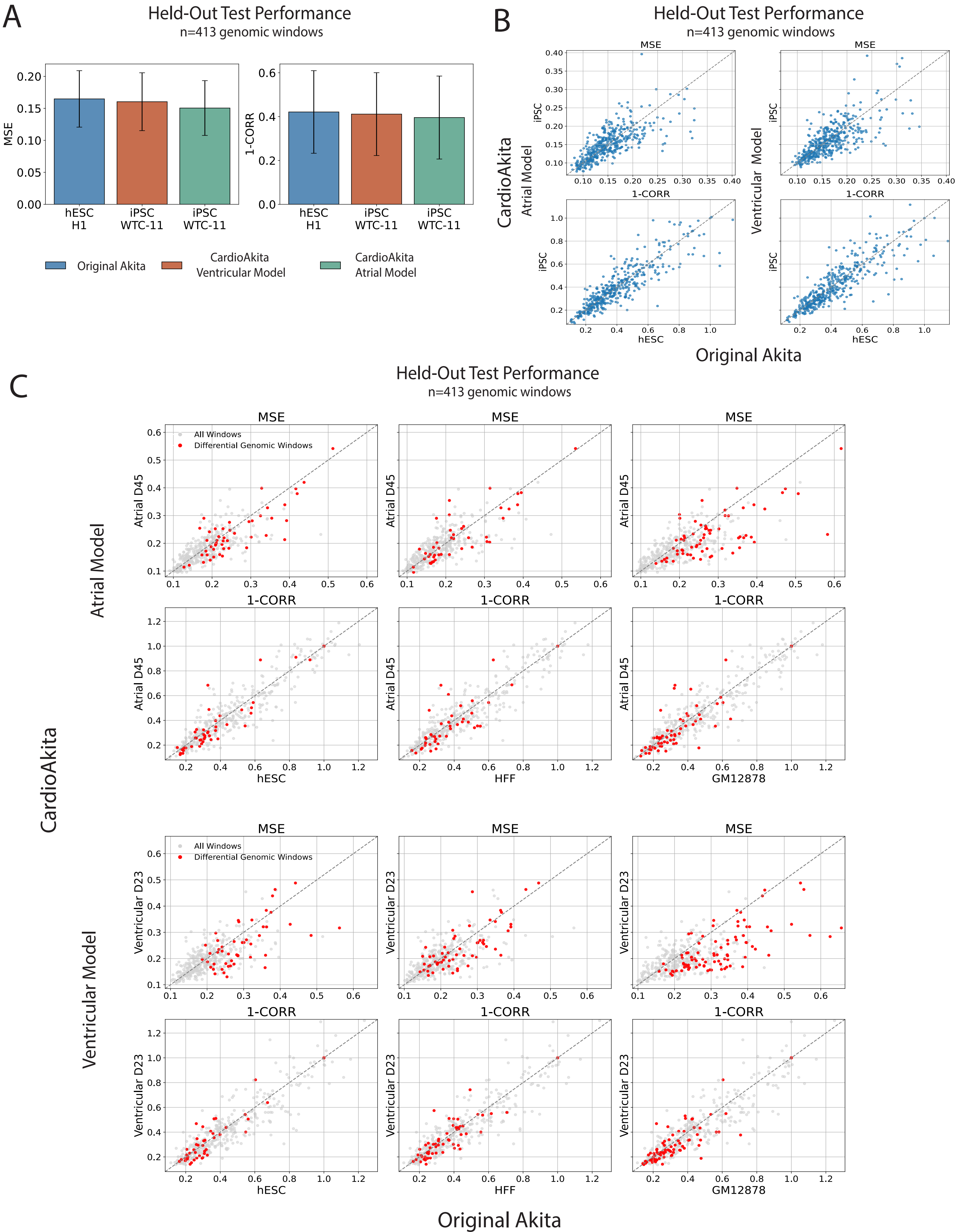

### Extended Data Figure 2

#### CardioAkita Disruption Scores:

##### Atrial Time Points:

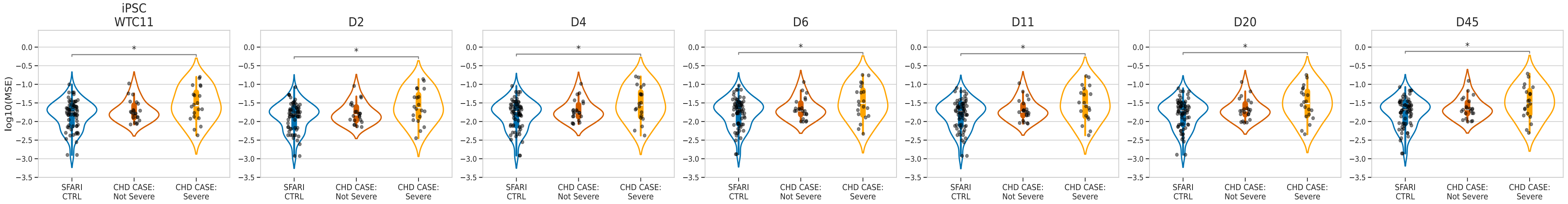


##### Ventricular Time Points:

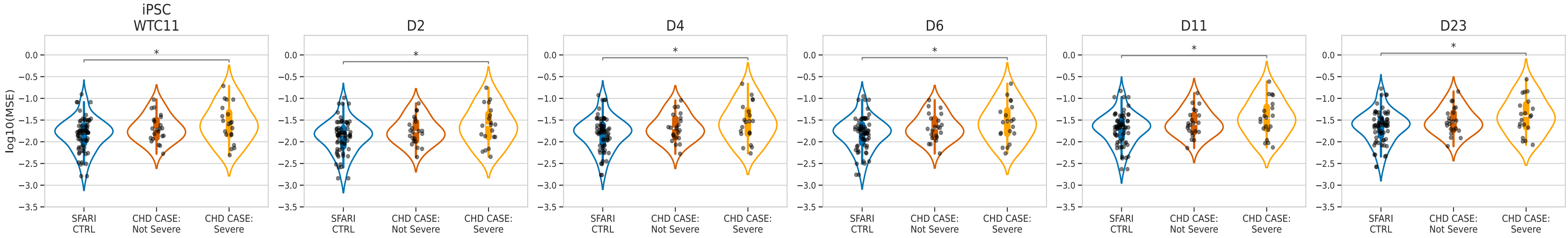

### Extended Data Figure 3

#### Supplementary Figure 3

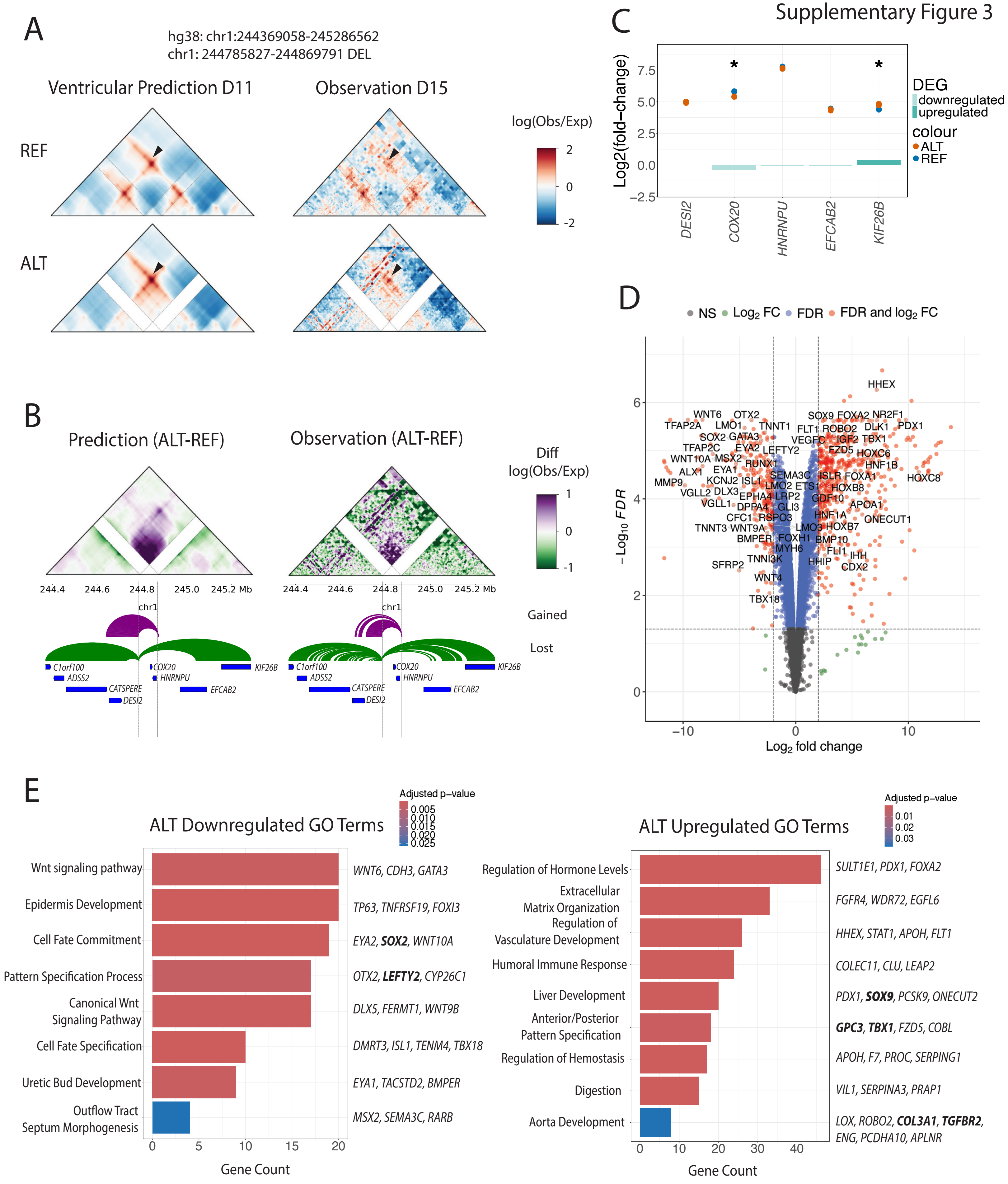

### Extended Data Figure 4

Supplementary Figure 4

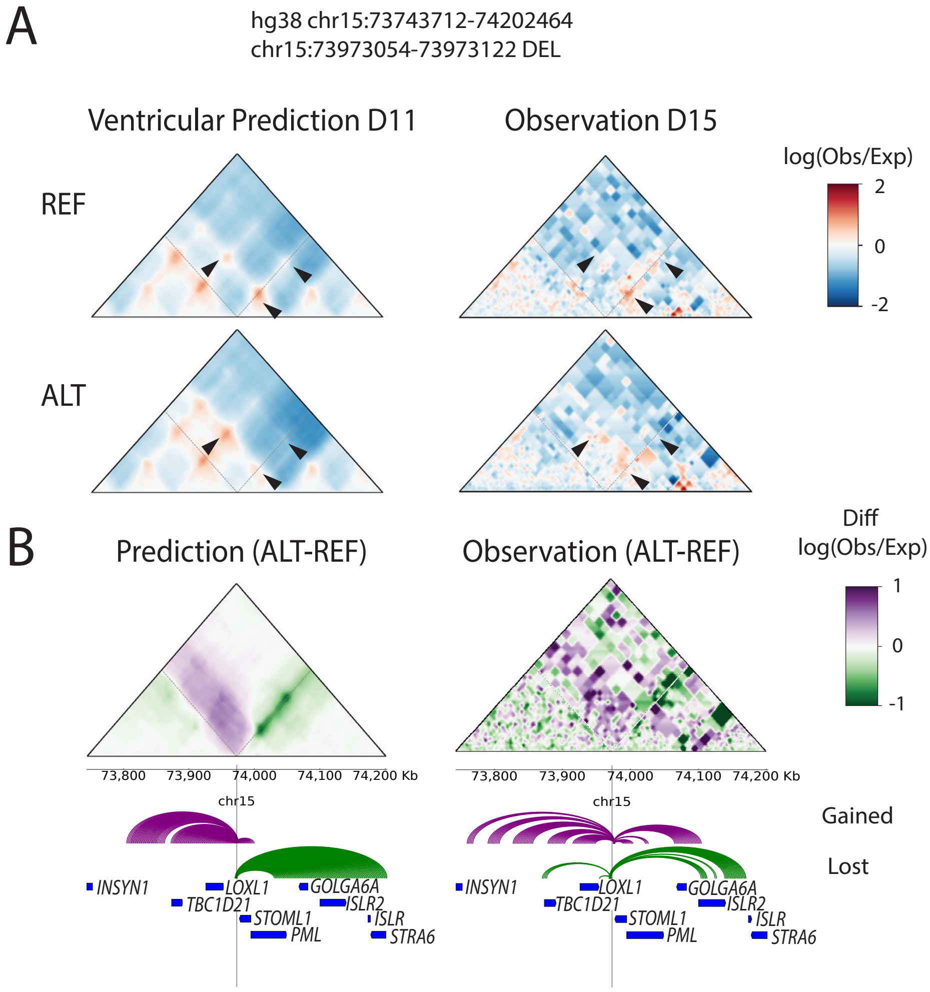
