## Extended Data Figure 5 for "Structural variants in human congenital heart disease disrupt distal genomic regulatory contacts of developmental genes"

A Heterozygous 58 kbp Deletion vs Ctrl

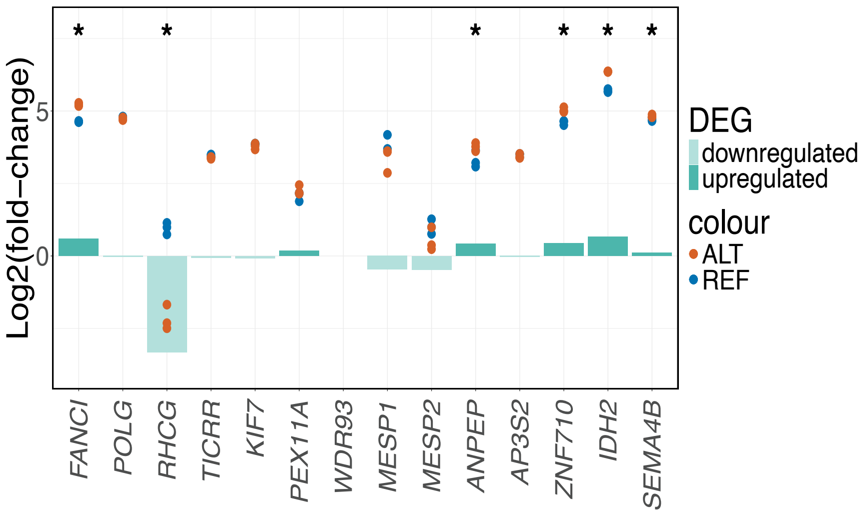


B Heterozygous 58kbp Deletion vs Ctrl

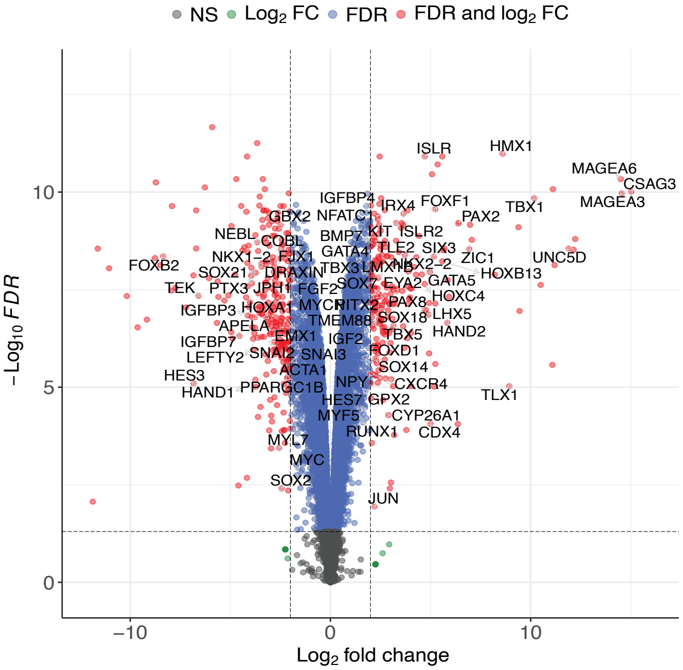


C

58 kbp Homozygous DEL and 58 kbp Heterozygous DEL Shared Upregulated Genes

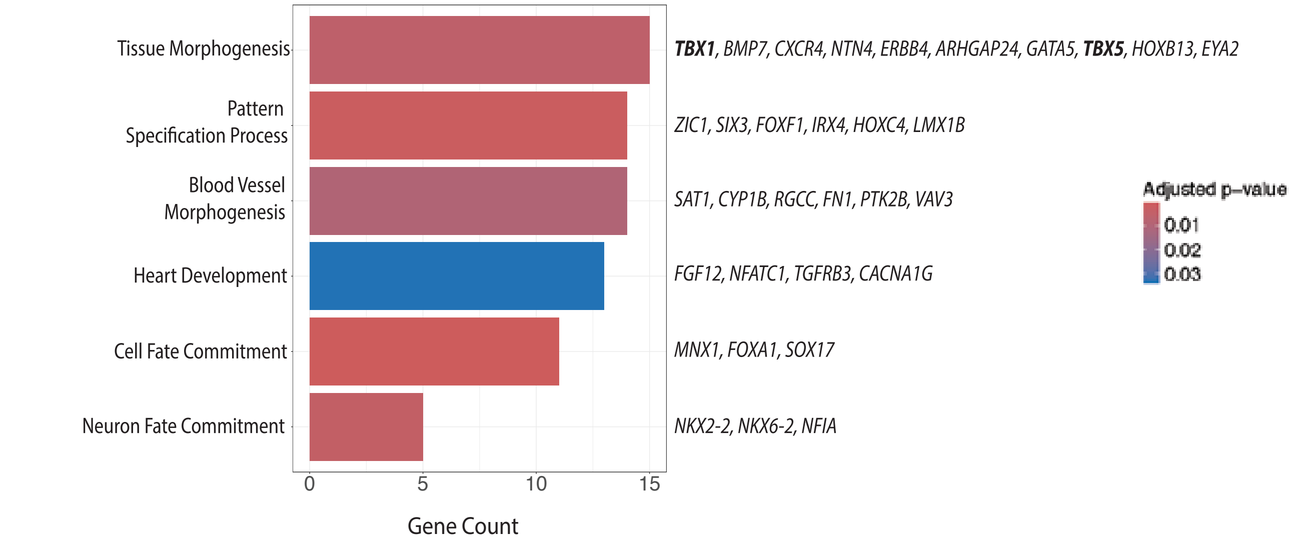


58 kbp Homozygous DEL and 58 kbp Heterozygous DEL Shared Downregulated Genes

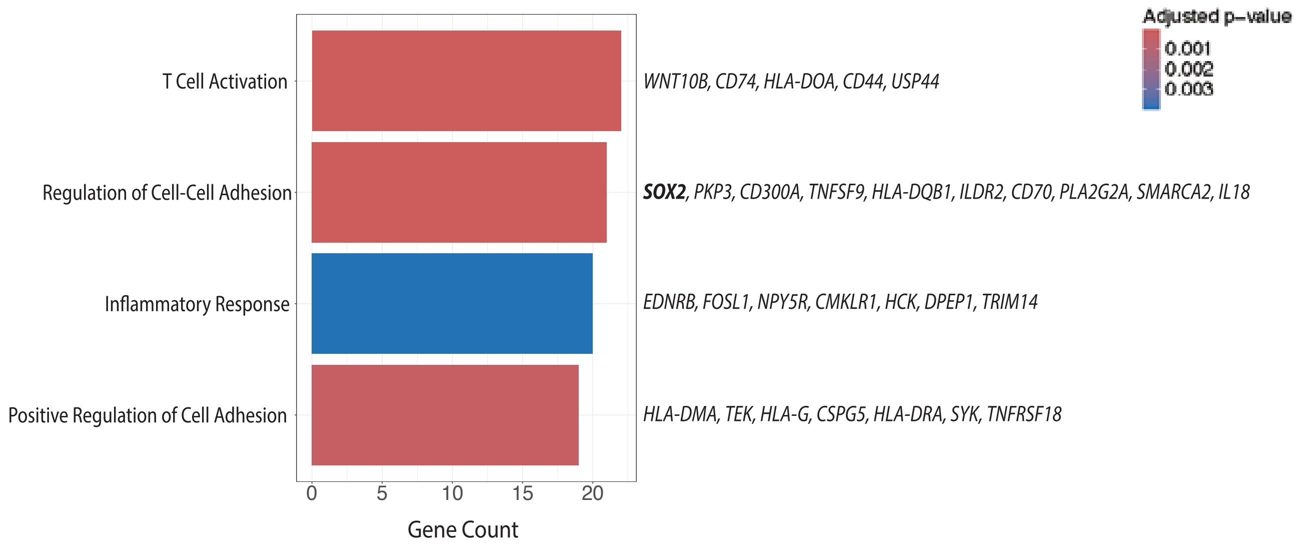
